# Structural insights into a novel family of integral membrane siderophore reductases

**DOI:** 10.1101/2021.01.28.428567

**Authors:** Inokentijs Josts, Katharina Veith, Vincent Normant, Isabelle J. Schalk, Henning Tidow

**Author notes:** European Molecular Biology Laboratory (EMBL), Hamburg Outstation, Hamburg, Germany. Corresponding authors: Inokentijs Josts, University of Hamburg, Department of Chemistry, Institute for Biochemistry and Molecular Biology, Luruper Chaussee 149, D-22761 Hamburg, Germany, Henning Tidow, University of Hamburg, Department of Chemistry, Institute for Biochemistry and Molecular Biology, Luruper Chaussee 149, D-22761 Hamburg, Germany.

## Abstract

Gram-negative bacteria take up the essential ion Fe^3+^ as ferric-siderophore complexes through their outer membrane using TonB-dependent transporters. However, the subsequent route through the inner membrane differs across many bacterial species and siderophore chemistries and is not understood in detail. Here, we report the crystal structure of the inner membrane protein FoxB (from *P. aeruginosa*) that is involved in Fe-siderophore uptake. The structure revealed a novel fold with two tightly-bound heme molecules. In combination with functional studies these results establish FoxB as an inner membrane reductase involved in the release of iron from ferrioxamine during Fe-siderophore uptake.

## Introduction

Most microbial communities rely on the bioavailability of iron for their survival. Due to its physico-chemical properties ferric iron exhibits low solubility at physiological pH, precipitating out as ferric oxides and hydroxides. The limited availability of this precious resource has led to severe competition amongst microbes to scavenge this extremely scarce micronutrient from the environment. Most bacterial species, as well as several fungi, secrete numerous small-molecule iron chelators, named siderophores, which bind ferric ion with extremely high affinity and selectivity, at the same time maintaining them in a soluble state. Once such a complex is formed, ferric-siderophores can be taken up into the cells. In Gram-negative bacteria, siderophores are generally taken up by TonB-dependent transporters (TBDTs) present in the bacterial outer membrane (OM). Translocation across the OM relies on the coupling of ferri-siderophore-bound TBDTs to the energizing complex in the inner bacterial membrane, comprised of ExbB/ExbD/TonB proteins. Generally, a single TBDT recognizes a subset of siderophores within the same chemical family.

The route of ferric-siderophores across the inner membrane (IM) is less straightforward and differs across many bacterial species and siderophore chemistries. Generally, once inside the periplasmic space, ferric-siderophore complexes are recognized by the dedicated periplasmic-binding proteins for delivery to IM transporters for uptake into the cytoplasm. The most common family of IM transporters involved in the uptake of ferric-siderophores are ATP-binding cassette (ABC) importers. Several ABC transporters implicated in siderophore transport have been identified and characterized to date (Schalk and Guillon 2013) (Rohrbach, Braun et al. 1995).

In order to assimilate the captured iron into their biological processes, the ferric-siderophore complex must be dissociated. Several mechanisms for iron release from the chelated siderophores have been postulated, which finally all lead to iron reduction, associated either with a siderophore hydrolysis (Miethke and Marahiel 2007), chemical modification of the siderophore such as acetylation (Hannauer, Barda et al. 2010) or proton-mediated iron release (Marshall, Stintzi et al. 2009) (Schroder, Johnson et al. 2003) (Miethke and Marahiel 2007). Hydrolysis of siderophores is carried out by dedicated esterases, which fragment the siderophore, lowering the stability of the interaction with iron and facilitating subsequent dissociation steps. To-date, several siderophore hydrolases and esterases have been identified, including Fes, IroD and IroE found in *E. coli* and *Salmonella* (Zhu, Valdebenito et al. 2005) (Lin, Fischbach et al. 2005), PfeE from *P. aeruginosa* (Perraud, Moynie et al. 2018) and Cee from *Campylobacter jejuni* (Zeng, Mo et al. 2013). Some of these proteins are cytoplasmic, whereas others are localized in the periplasm.

The bacterial cell compartment of iron reduction and release from the siderophore complex also differs based on the microbe and siderophore. Several siderophore reductases have been identified in various bacteria and fungi. For example, *E. coli* have FhuF, a Fe-S cluster containing protein that targets ferric complexes with coprogen, ferrichrome and ferrioxamine B (Matzanke, Anemuller et al. 2004). Additionally, YqjH, another siderophore reductase found in *E. coli,* as well as its homologue ViuB from *Vibrio cholera,* use NADPH-FAD co-factors to reduce iron in ferric-triscatecholates complexes such as enterobactin, aerobactin, vibriobactin and ferric dicitrate (Miethke, Hou et al. 2011). In Gram-positive mycobacteria, IrtAB, a fusion of an ABC transporter and a reductase is responsible for both the uptake and subsequent reduction of several myco-bacterial siderophores (Arnold, Weber et al. 2020). All these siderophore reductases function in the cytoplasm after the translocation of the ferric-chelate across the bacterial IM.

In stark contrast, certain siderophores rely on iron release within the periplasmic space. For example, in *P. aeruginosa* pyoverdine and citrate siderophores dissociate their iron in the periplasm for subsequent uptake via dedicated Fe transporter, implying the existence of siderophore-specific periplasmic reductases (Greenwald, Hoegy et al. 2007) (Ganne, Brillet et al. 2017) (Marshall, Stintzi et al. 2009). Recent studies have identified the four proteins FpvG, FpvH, FpvJ and FpvJ, which co-occur in the pyoverdine uptake operon are responsible for the reduction of iron in the ferric-pyoverdine complex (Ganne, Brillet et al. 2017). The inner membrane FpvG is the reductase, but it needs the two other inner membrane proteins FpvH and FpvI and the periplasmic FpvJ protein for its full activity.

*Pseudomonas aeruginosa* is a ubiquitous Gram-negative bacterium that has emerged as an opportunistic human pathogen with significant clinical implications. It is one of the leading causes in high mortality rates arising from nosocomial infections. It uses two major siderophores, namely pyoverdine and pyochelin, to obtain iron from the environment. At the same time, it can also utilize foreign siderophores (exosiderophores) as part of its highly adaptable life-style. One such exosiderophore family consists of hydroxamate polydentates named ferrioxamines produced by members of *Streptomyces* family and several fungi. *P. aeruginosa,* like numerous Gram-negative bacteria, is able to take up ferrioxamines via a dedicated TBDT FoxA in an act of siderophore piracy (Llamas, Sparrius et al. 2006) (Josts, Veith et al. 2019). Interestingly, our recent studies have suggested that ferrioxamine B but not ferrioxamine E (nocardamine) uses an additional, unidentified TBDT for its uptake into *P. aeruginosa* (Normant, Josts et al. 2020). Generally, Gram-negative bacteria use ABC transporters *fhuDBC* or *hmuUVT,* in a species-specific manner to transport hydroxamate siderophores across the IM (Cuiv, Keogh et al. 2008) (Kingsley, Reissbrodt et al. 1999). However, neither of these uptake systems have been described or characterized in *P. aeruginosa*. Here, we report on the crystal structure of FoxB, which belongs to a large, poorly understood family of inner membrane oxidoreductases associated with siderophore uptake and processing. The protein possesses a novel fold with the transmembrane domain harboring di-hemes indicating a role as inner membrane reductase involved in Fe-siderophore uptake.

## Results

### Overall structure of FoxB

Our previous biochemical and structural studies have implicated the TBDT FoxA in uptake of ferrioxamines B and E across the OM in *P. aeruginosa*. The *foxA* gene shares its operon with genes encoding for sigma and anti-sigma factors *foxI* and *foxR,* respectively, which regulate the operon expression through siderophore-dependent signaling cascades involving the signaling domain of FoxA (Fig. 1a). The operon possesses an additional gene, coding for an uncharacterized IM protein, FoxB, belonging to the conserved family of iron-regulated membrane proteins, termed the PepSY_TM family (COG3182). FoxB and its distant orthologues have been implicated in siderophore uptake but with little functional understanding of their activity (Cuiv, Keogh et al. 2007). Recent studies with a paralogous protein FpvG, found in the pyoverdine operon, have concluded FpvG to be an integral membrane siderophore reductase of ferric pyoverdine (Ganne, Brillet et al. 2017). Therefore, we reasoned that FoxB could also act as a ferrioxamine reductase. To gain insights into the proposed reductase function of FoxB, we have determined the crystal structure of FoxB using X-ray crystallography.

**Figure 1.**
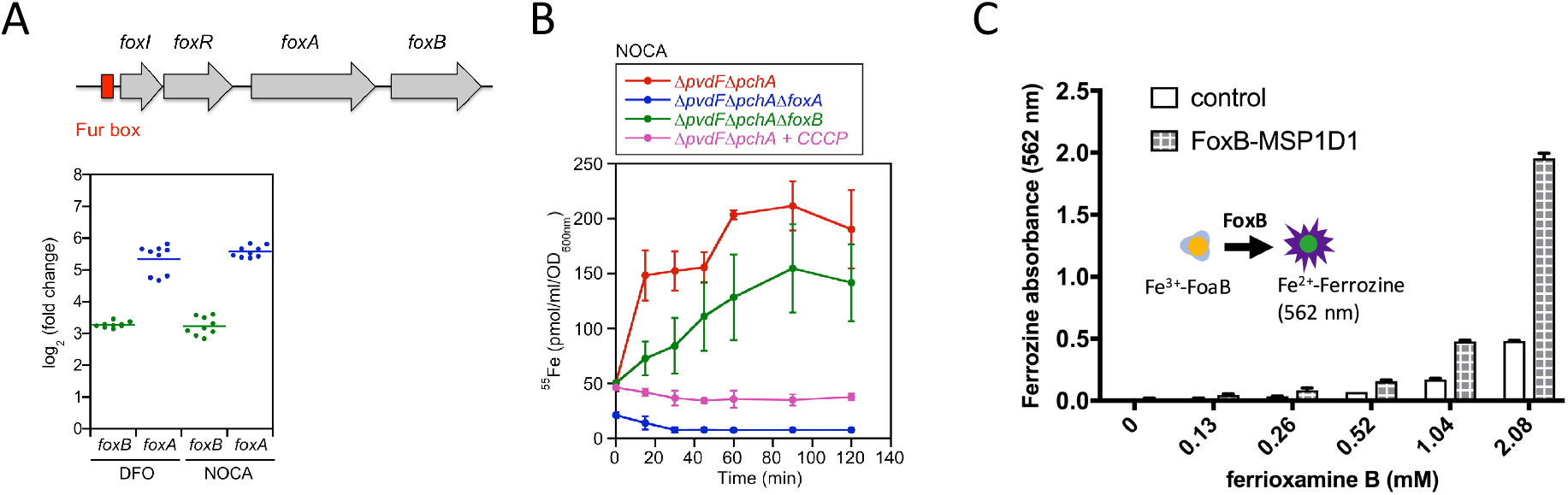
Functional characterization of *foxB*. A) Expression of *foxB* and *foxA* in presence of DFO or NOCA in minimal medium. Top panel: Organization of *fox* genes in *P. aeruginosa* genome. *foxA* encode for TBDT transporter localize at the outer membrane, *foxI* encode for sigma factor, *foxR* encode for anti-sigma factor localize at the inner membrane and *foxB* encode for protein localize at the inner membrane. A fur box is present upstream *foxI* gene. Bottom panel: PAO1 WT was grown in CAA medium overnight and, then resuspended in fresh CAA medium at OD_600nm_ = 0.1 and treated or not with 10 μM DFO or 10 μM NOCA for 8 hours. Total RNA was extracted and retro-transcribed in cDNA. Expression of genes was normalized to the *uvrD* reference gene in each condition. The expression of *foxA* and *foxB* was measured in DFO or NOCA condition and related to untreated condition. Results were represented in log_2_ fold change expression. Graphical represent the result of three biological experiments each performed in triplicate. The bars represent the means. B) Transport of ^55^Fe-NOCA in PVD and PCH biosynthesis deficient cells expressing or not *foxA* or *foxB*. PAO1 *ΔpvdFΔpchA*, *ΔpvdFΔpchAΔfoxA* and *ΔpvdFΔpchAΔfoxB* were grown in CAA medium with 10 μM NOCA to induce *foxA* and *foxB* expression overnight at 30°C. Then, cells were washed with 50mM Tris-HCl (pH 8.0). The transport assay was started by the addition of 500 nM ^55^Fe-NOCA to *ΔpvdFΔpchA* (red curve), *ΔpvdFΔpchAΔfoxA* (blue curve) and *ΔpvdFΔpchAΔfoxB* (green curve). The *ΔpvdFΔpchA* strain was also treated with 200 μM CCCP protonophore (pink curve) as a control in presence of ^55^Fe-NOCA. At each indicated point (0 to 120 minutes), samples were centrifuged, and the radioactivity retained in the cell pellets were monitored. Data represent the mean of ^55^Fe concentration in pmol/ml/OD_600nm_ ± SD of three independent experiments. C) Ferrioxamine B reduction assay using ferrozine absorption to monitor Fe^2+^ release from the siderophore. Absorbance was measured at 562 nm for a range of ferrioxamine concentrations and FoxB in MSP1D1 (3 μM) and 15 mM DTT. Plots represent an average of three independent assays with standard error shown.

Upon over-expression and purification of FoxB from *E. coli* membranes it became evident that the protein possesses heme groups with a typical UV-Vis spectrum having a Soret band and a broad Q-band, characteristic of heme-bound proteins. Reducing treatment of FoxB with sodium dithionite resulted in the shift of the Soret peak from 414 nm to 429 nm, along with a sharp appearance of distinct peaks in the Q-band region (Suppl. Fig. S1a). Oxidation of FoxB with potassium ferricyanide reverted the spectrum to its original state, indicating that FoxB purifies in an oxidized state. Reducing activity of FoxB and subsequent iron dissociation from ferrioxamine B was followed using the colorimetric ferrozine assay, which detects free Fe^2+^ ions in solution. Purified FoxB had no detectable reductive capacity in any detergents. However, after reconstitution into MSP1D1 nanodiscs, the reductive activity of FoxB could be measured, using a range of Fe^3+^-ferrioxamine B concentrations (Fig. 1c). Ferrozine forms a complex with Fe^2+^ resulting in absorbance at 562 nm and a colour change of solution. The increase in absorbance at 562 nm could be followed over time in the presence of FoxB, DTT and Fe^3+^-ferrioxamine B, indicating that Fe^2+^ ions were formed by the reduction of the ferric sideophore. We determined the crystal structure of FoxB using single-anomalous dispersion (SAD) in combination with molecular replacement. Initial SAD phases based on the calculated selenium and iron sites (Suppl. Fig. S2) allowed the building of approximately 70% of the backbone revealing the overall fold of FoxB and placement of heme groups. Some sequence assignment could be made based on the positions of Se atoms, however due to the low number of methionine residues (5 total out of 382 residues) and low resolution of the datasets further side chain tracing could not be completed. We then submitted our sequence to the CASP14 (Critical Assessment of Structure Prediction) competition (target T1058) and used the model predicted by the best-performing group (later identified as AlphaFold2 / DeepMind) (Callaway 2020) for molecular replacement. With this model we obtained a clear MR solution, which could then be used for MR-SAD resulting in a good electron density map for entire protein (Suppl. Fig. S3). The structure consists of a four transmembrane (TM) helical bundle capped at the periplasm by two PepSY (peptidase propeptide and YpeB domain / Pfam:PF03413) domains (Fig. 2). The cytoplasmic residues forming a loop between TM2 and TM3 (residues 172 -188) were absent from the electron density and omitted from the final model. The TM bundle wraps around two b-type hemes coordinated non-covalently by highly conserved His residues protruding from the TM domains (Fig. 2 / Suppl. Fig. S4). The iron in both heme groups is in the octahedral coordination state. Both hemes are found at the edges of the lipid bilayers with propionate groups protruding towards the solvent environment. The positioning of the TM bundle in the lipid bilayer was analyzed by the OPM web server (Lomize, Pogozheva et al. 2012) and suggests a tilt angle of 20° with a hydrophobic thickness of 30.4 Å (Suppl. Fig. S5). TM3 is kinked in the middle by a non-conserved stretch of G_201_-G_202_ residues, splitting it into 2 TM helices, TM3a and TM3b. The kink causes TM3a to bend by 66° perpendicular to the lipid bilayer plane and positions H198 to act as an apical ligand coordinating Fe within the cytoplasmic heme group.

**Figure 2.**
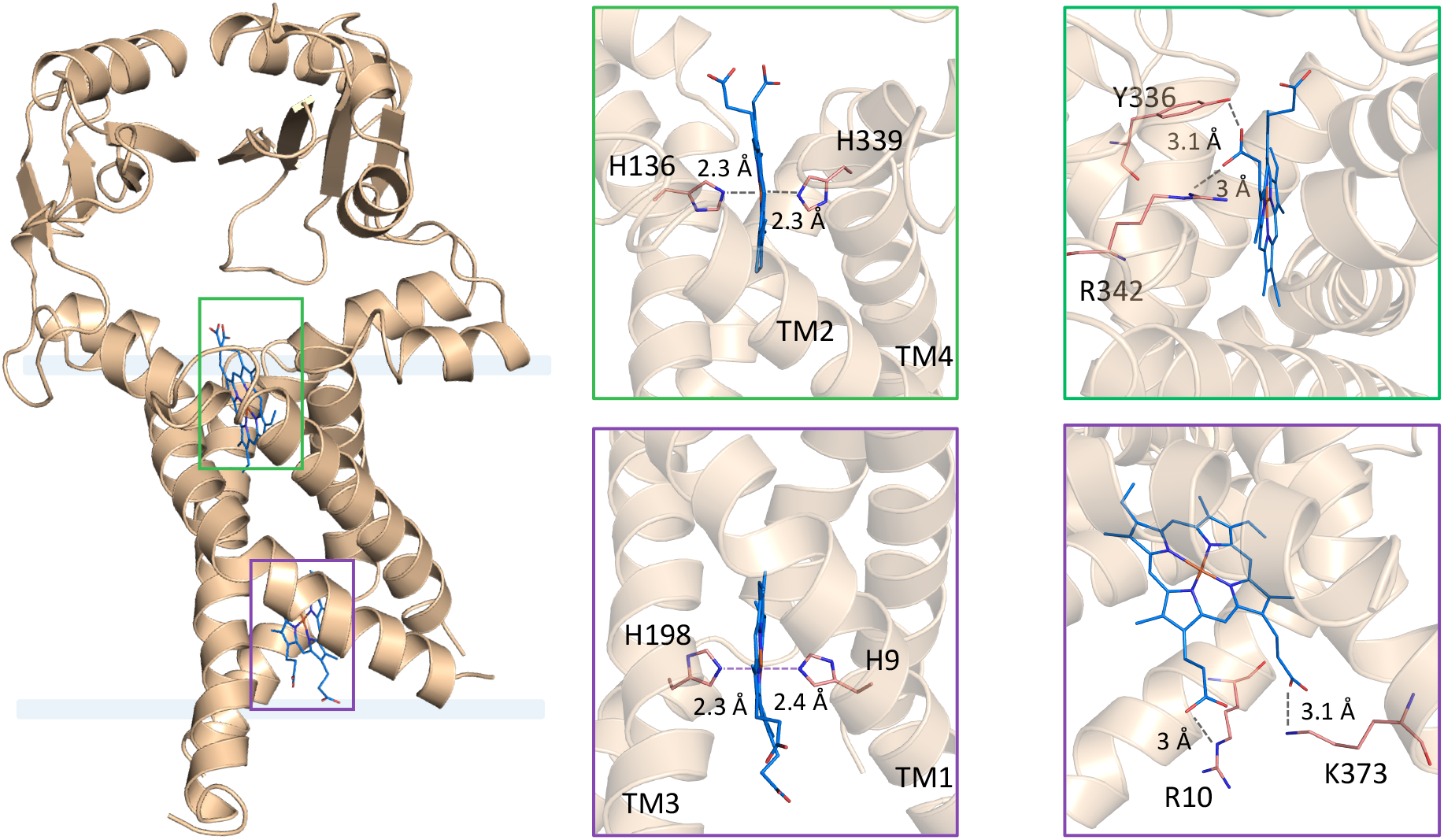
Overall structure of FoxB. A) FoxB structure reveals a 4-helix TM-fold topped with periplasmic PepSY domains. The membrane-spanning region is indicated and both heme molecules are boxed. B) Di-heme coordination by the TM domain of FoxB. All four transmembrane helices contribute to heme binding. Both hemes are found at the edges of the lipid bilayers with propionate groups protruding towards the solvent environment. Coordinating residues towards the Fe atoms or propionate groups are indicated for both hemes in two orientations.

Both propionate groups of the cytoplasmic heme coordinate the conserved residues R10 and K373 through electrostatic or H-bond forces. At the periplasmic side a single heme propionate group is within H-bonding distance of a conserved R342 and a non-conserved Y336 (Fig. 2). The periplasmic PepSY domains form extensive inter-domain contacts through β-augmentation between the β-sheets of individual domains as well as additional side-chain H-bond contacts from the adjacent α-helices (D260 and S55) (Fig. 3). The buried surface area for the interdomain complex is 479.3 Å^2^. Both PepSY domains enclose a large, solvent-accessible cavity located above the TM domain (Suppl. Fig. S6). This cavity is rich in aromatic amino acids along with a cluster of charged residues in PepSY domain 1. The cluster of negatively charged residues is located on the periplasmic surface of domain 1 (residues 33-117) and a loop from the second PepSY domain (residues 297-300) (Suppl. Fig. S7a).

**Figure 3.**
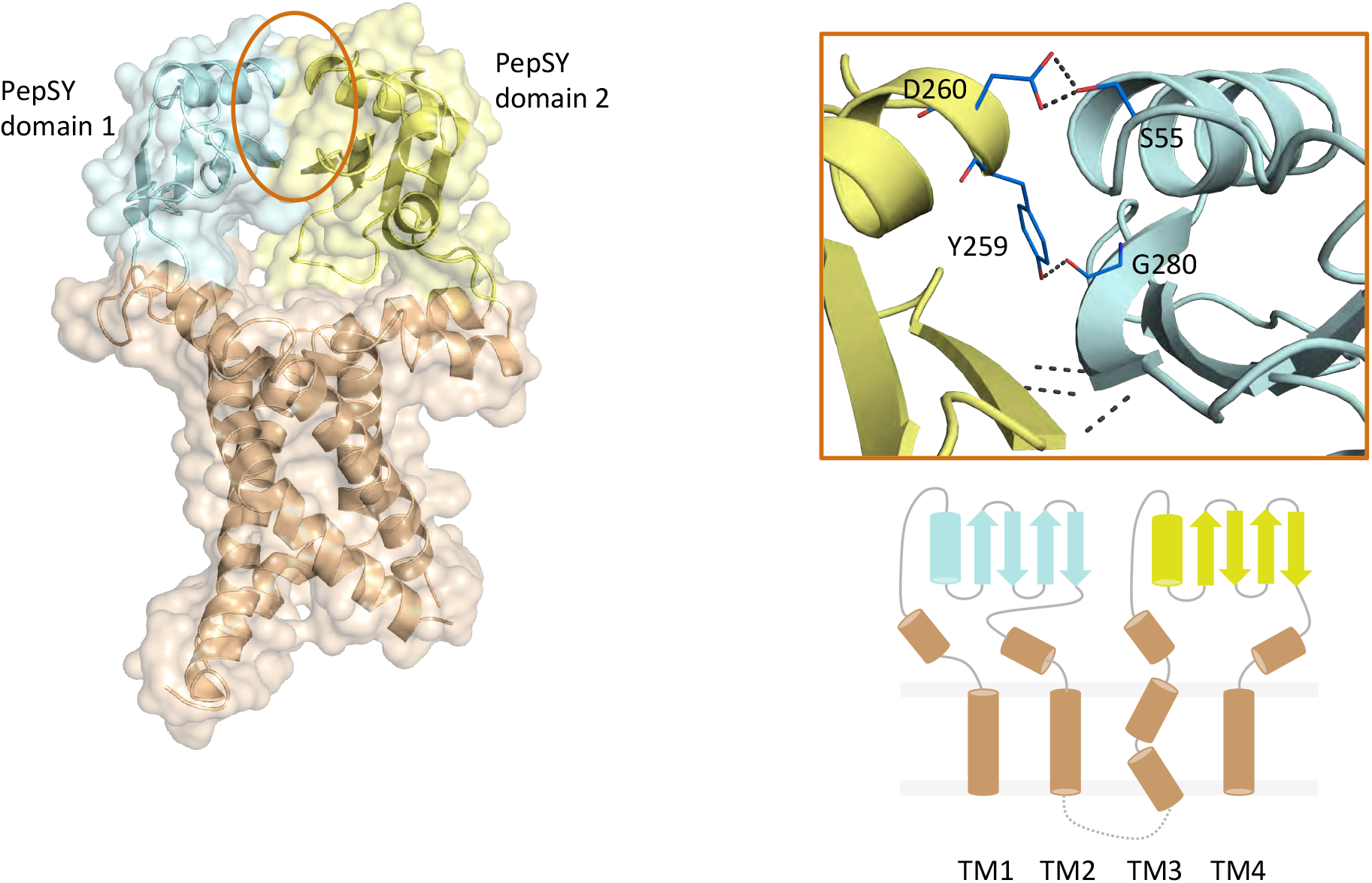
Periplasmic PepSY domains within FoxB. A) Two PepSY domains (residues 46-118 and 239-325, respectively) form the periplasmic part of the protein and interact via several hydrogen bonds (shown in B). Overall secondary structure of FoxB is also shown.

We could identify two Zn ions in one of the molecules of FoxB (Zn_A1_ and Zn_A2_) as well a single Zn atom within the second FoxB molecule (Zn_B_). The crystallization conditions contain 100mM ZnCl_2_, which we believe is the source for bound Zn ions. No crystals of FoxB could be obtained without Zn supplementation. Zn_A1_ and Zn_A2_ are coordinated by a tetrad of His residues, which form a small tunnel out of the periplasmic cavity. The position of Zn_A1_ and Zn_B_ are equivalent but not identical, both ions being in the vicinity of the propionate groups of the periplasmic heme group (Suppl. Fig. S7b).

DALI search suggest a very distant structural similarity of the FoxB TM domains to the *E. coli* superoxide:ubiquinone oxidase (pdb:5oc0) and fumarate reductase (pdb:1e7p) with a root mean square deviation of 5.6 and 5.5 Å, respectively (Suppl. Fig. S8a). Both of these are heme-containing membrane proteins involved in redox reactions. Analysis of the PepSY domains from FoxB against the available structures of PepSY-containing proteins suggests a highly conserved fold (Suppl. Fig. S8b). The function of these proteins is currently poorly understood with experimental evidence pointing towards inhibition of peptidase activity through protein-protein interactions. However, these proteins are believed to not possess any TM domains and regulate specific peptidases as stand-alone entities (Yeats, Rawlings et al. 2004).

### DFO and NOCA induce *foxB* and *foxA* expression

A Fur box is present upstream of *foxI* gene (sigma factor) and represses the transcription of *fox* genes in the presence of iron. In iron deficient conditions, *foxA* expression is induced (Llamas, Sparrius et al. 2006); this expression is enhanced when the exosiderophores DFO or NOCA are present in the *P. aeruginosa* environment (Llamas, Sparrius et al. 2006) (Normant, Josts et al. 2020). The upregulation of *foxA* transcription and expression is dependent of the presence of FoxI (Llamas, Sparrius et al. 2006). It has been shown that DFO induces the self-cleavage of the anti-sigma factor FoxR that certainly leads to the release of the sigma factor FoxI (Bastiaansen, Otero-Asman et al. 2015).

To check whether *foxB* transcription is induced in the presence of DFO or NOCA (as it is the case for *foxA*), *P. aeruginosa* PAO1 strain was grown in CAA iron deficient medium with and without 10 μM DFO or NOCA. After 8 hours, total RNA was extracted and *foxA* and *foxB* transcription were measured by RT-qPCR. As previously described (Normant, Josts et al. 2020), an increase in *foxA* transcription in the presence of both DFO and NOCA was observed (log_2_ fold change increase of 5.34 ± 0.45 for DFO and 5.57 ± 0.18 for NOCA) (Fig. 1a). In parallel, *foxB* transcription was also increased in the presence of DFO (log_2_ fold change increased 3.28 ± 0.09) and NOCA (log_2_ fold change increased 3.23 ± 0.28), demonstrating that transcription of both *foxA* as well as *foxB* genes is regulated by the presence of DFO and NOCA.

### *foxB* deletion affects iron acquisition by NOCA

We have previously shown that iron acquisition by NOCA was completely dependent on FoxA outer membrane transporter expression, but iron uptake by DFO was only partially dependent on this transporter (Normant, Josts et al. 2020). To investigate the role of FoxB in these two iron acquisition pathways, ^55^Fe uptake was monitored in the presence of these two exosiderophores in a PVD and PCH deficient *P. aeruginosa* strain (Δ*pvdF*Δ*pchA*) and its corresponding *foxA* and *foxB* deletion mutants (Δ*pvdF*Δ*pchA*Δ*foxA* and Δ*pvdF*Δ*pchA*Δ*foxB*) (Fig. 1b). Strains unable to produce pyoverdine and pyochelin were used to avoid any ^55^Fe uptake by these two siderophores. These strains were grown in iron restricted CAA medium with 10 μM DFO or NOCA to induce the expression of *foxA* and *foxB* and the uptake assays were initiated by addition of 500 nM ^55^Fe-DFO or ^55^Fe-NOCA.

Similar rates as previously described for ^55^Fe uptake in the presence of DFO and NOCA in *ΔpvdFΔpchA* cells were observed (Normant, Josts et al. 2020). In the *foxA* deletion mutant (*ΔpvdFΔpchAΔfoxA*), transport of ^55^Fe-NOCA was totally abolished and partially affected for ^55^Fe-DFO as previously described (Fig. 1b): ferri-NOCA is transported across the outer membrane only by FoxA while ferri-DFO is transported by FoxA and at least another TonB-dependent transporter (Normant, Josts et al. 2020). *foxB* deletion had no effect on ^55^Fe assimilation in the presence of DFO, but partially affected iron uptake by NOCA: 37 % inhibition after 60 minutes and around 26 % after 120 minutes (Fig. 1b). The absence of *foxB* expression altered iron acquisition by NOCA but not the one by DFO, indicating that FoxB plays a role in iron assimilation by NOCA but is not essential for the DFO iron uptake pathways. FoxB can probably be replaced by another protein that has a redundant function in ferri-NOCA as well as in ferri-DFO uptake pathways.

### Members of COG3182 family are widely distributed across the bacterial kingdom

Search for FoxB paralogues in the *P. aeruginosa* PAO1 genome revealed the existence of up to five additional operons associated with siderophore or metal ion uptake bearing COG3182 gene members. Each operon also has a dedicated TBDT adjacent to the COG3182 gene. The operons and their associated gene members are outlined in Suppl. Fig. S9. They include the pyochelin operon, putative mycobactin operon and three additional “orphan” operons with no known siderophore substrates (ferrioxamine and pyoverdine operons are excluded).

Members of COG3182 family are widespread across the bacterial kingdom including proteobacteria, firmicutes, bacteroides and cyanobacteria. In Gram-negative bacteria, COG3182 members are often in genomic proximity of a TBDT associated with uptake of siderophores and metal ions as judged by the COGNAT analysis. In several instances we find gene fusions between COG3182 members and sulfite reductases.

## Discussion

Siderophores act to deliver iron, an essential micronutrient, inside the cells of bacteria and fungi. Uptake and translocation of siderophores have received significant scientific attention, yet the precise mechanisms of iron release from these chelated complexes still remain poorly understood. In this study, we report on the structure of FoxB, which belongs to a large, uncharacterized family of integral membrane oxidoreductases that assist in the assimilation of iron from the captured ferric-siderophore complexes in the periplasm through reduction and release of iron from siderophores.

Based on the structure of FoxB, we propose a through-space electron pathway involving the heme molecule on the cytoplasmic side of the TM-region, two conserved Trp residues (W157/W206) located in TM2/3a, the periplasmic heme as well as two Zn^2+^ ions that may occupy/substitute the Fe^3+^-siderophore binding site. All residues involved are aligned within a distance of 5 Å (Fig. 4). Initial studies into the possible function of FoxB have shown that it was able to substitute the *rhtX* deletion in uptake of schizokinen, another hydroxamate siderophore in *Sinorhizobium meliloti* (Cuiv, Clarke et al. 2004). RhtX and its *Pseudomonas* homologue FptX are members of the major facilitator family of transporters and their roles in siderophore transport are established. We found up to five additional genes (FoxB and FpvG excluded) belonging to the COG3182 family in *P. aeruginosa* PAO1 strain.

**Figure 4.**
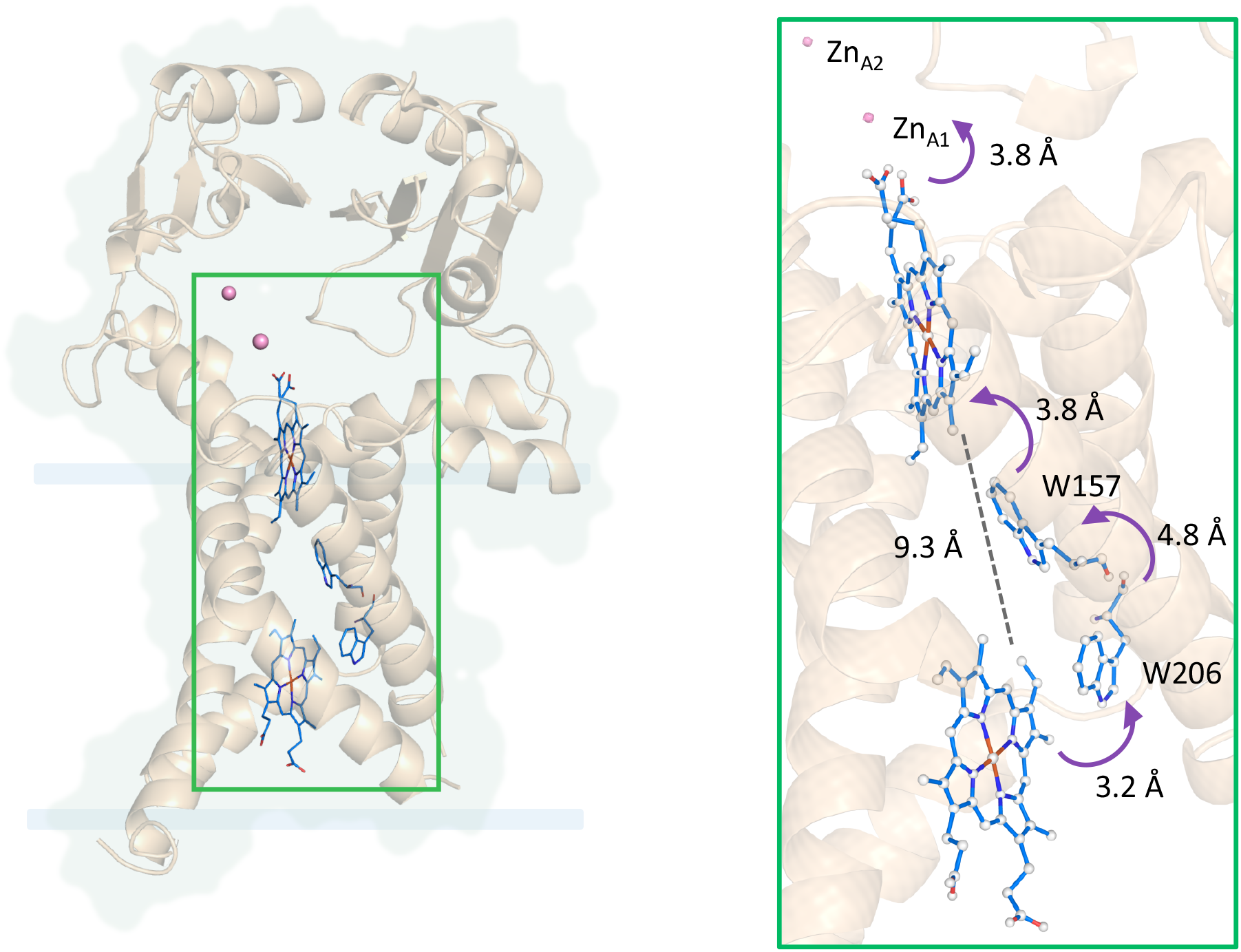
Potential electron pathway in FoxB reductase. The proposed through-space electron pathway comprises the heme molecule on the cytoplasmic side of the TM-region, two highly conserved Trp residues (W157/W206) located in TM2/3a, the periplasmic heme as well as two Zn^2+^ ions that may occupy/substitute the Fe^3+^-siderophore binding site. All residues involved are aligned within a distance of 5 Å.

Deletion of *foxB* leads to reduced uptake of iron from NOCA but not ferrioxamine B in *P. aeruginosa*. This result somewhat mirrors our previous study of the TBDT transporter FoxA present in the same operon as FoxB (Josts, Veith et al. 2019). FoxA is able to recognize both ferrioxamine B and E with same propensity, however deletion of the *foxA* gene does not abolish ferrioxamine B uptake (Normant, Josts et al. 2020). Redundancy amongst TBDT towards siderophore uptake is common. The ability of FoxB to assist in the uptake of schizokinen, a hydroxamate siderophore, in *S. meliloti* could also indicate a degree of promiscuity of FoxB-like proteins in *P. aeruginosa*. An alternative explanation is that ferrioxamine B could be taken up directly into the cytoplasm by an uncharacterized IM transporter. In *E. coli,* hydroxamate siderophores such as ferrioxamine B are taken up via the FhuBCD ABC transporter, which is not present in *P. aeruginosa.* However, an operon consisting of genes PA2912-2914 encodes for a putative ABC transporter with distant homology to HmuUVT in *P. aeruginosa.* This protein complex has been implicated in heme and hydroxamate siderophore uptake in *S. meliloti* (Cuiv, Keogh et al. 2008). Studies investigating the role of FpvG, a pyoverdine IM reductase in the COG3182 family, have also shown that *fpvG* mutants reduce ^55^Fe uptake by pyoverdine in *P. aeruginosa* by approximately 30-50%. It is interesting that FpvG seemingly functions as part of a much larger membrane complex including FpvH, FpvI and FpvJ (Ganne, Brillet et al. 2017), while FoxB seems to function alone. The pyoverdine operon also possesses an ABC transporter *fvpCDEF,* which is implicated in the uptake of iron released from pyoverdine by FpvGH (Ganne, Brillet et al. 2017). Since deletion of this transporter reduces uptake by 40-50%, it is possible that iron finds its way into the cells via a different IM transporter. The function of ferrous iron transporter FeoB in the acquisition of iron from ferric-citrate has been demonstrated (Marshall, Stintzi et al. 2009), however it had no role in the transport of iron derived from ferrioxamine B or pyoverdine. Therefore, we cannot rule out the contribution of another IM protein in the translocation of iron ions inside *P. aeruginosa* cells.

## Materials and Methods

### Chemicals

Ferrioxamine B (DFO) and the protonophore carbonyl cyanide *m*-chlorophenylhydrazone (CCCP) were purchased from Sigma-Aldrich, ^55^FeCl_3_ was purchased from Perkin Elmer. NOCA was purified as previously described (Meyer and Abdallah 1980).

### Bacterial strains and growth conditions

The *P. aeruginosa* PAO1 strains used in this study are listed in Table S2. Bacteria were initially grown in LB medium overnight at 30 °C. Next, bacteria were washed and resuspended in iron-deficient CAA (casamino acid) medium containing 5 g l^−1^ low-iron CAA (Difco), 1.46 g l^−1^ K_2_HPO_4_ 3H_2_O and 0.25 g l^−1^ MgSO_4_ 7H_2_O and were grown overnight at 30 °C.

### Plasmid and strain constructions

For pEXG2*foxB* construction a 1449 bp insert containing flanking sequences of *foxB* gene (736 bp upstream and 715 bp downstream *foxB* gene) was amplified by PCR from genomic DNA of *P. aeruginosa* PAO1 using the specific primers listed in Table S4 with High-Fidelity DNA polymerase (Thermo-Fisher Scientific). Primers were designed to generate XhoI and HindIII restriction sites at 5’end and 3’end of the PCR product. The PCR product was cloned into pEXG2 vector using XhoI and HindIII restrictions sites. Ligation was performed using T4 DNA ligase (Thermo-Fisher Scientific). pEXG2*foxB* plasmid was transformed in *E. coli* strain TOP10 (Invitrogen). Mutation in the chromosomal genome of *P. aeruginosa ΔpvdFΔpchA* was generated like previously described (Perraud, Cantero et al. 2020) by triparental mating. Mutants were selected and verified by PCR and sequencing.

### RNA extractions

Bacteria were grown overnight in CAA medium. Afterwards, bacteria were pelleted, washed and resuspended at OD_600 nm_ = 0.1 in fresh CAA medium with or without 10 μM NOCA or DFO. Cells were cultivated at 30 °C for 8 hours. Then, 2.5 × 10^8^ cells were mixed with two volumes of RNAprotect Bacteria Reagent (Qiagen). Samples were lysed in Tris-EDTA (pH 8.0) containing 15 mg/ml lysozyme (Sigma-Aldrich) for 15 minutes at 25 °C. Lysates were homogenized with QIAshredder kit (Qiagen) and total RNAs were extracted with RNeasy mini kit (Qiagen). After treatment with DNase (RNase-Free DNase Set, Qiagen), RNAs were purified with an RNeasy Mini Elute cleanup kit (Qiagen).

### RT-qPCR analysis

RNA (1 μg) was reversed-transcribed with High Capacity RNA-to-cDNA Kit (Applied Biosystems), in accordance with the manufacturer’s instructions. The expression of genes was measured in a SpetOne Plus Instrument (Applied Biosystems), with Power Sybr Green PCR Master Mix (Applied Biosystems) and the appropriate primers (Table S3). The *uvrD* expression was used as an internal control. For a given gene in each strain, the transcript levels were normalized with respect to those for *uvrD* and were expressed as a base two logarithms of the ratio (fold-change) relative to the reference conditions.

### Iron uptake

Siderophore-^55^Fe complexes were prepared as previously described (Hoegy and Schalk 2014) with a ratio 20:1 for siderophore:iron, in 50 mM Tris-HCl (pH 8.0). After overnight growing in CAA medium, cells were washed and resuspended in fresh CAA medium at OD_600 nm_ of 0.1 in the absence of 10 μM NOCA or DFO to induce *foxA* and *foxB* expression, and grown overnight at 30 °C. Afterwards, bacteria were washed with 50 mM Tris-HCl (pH 8.0), and diluted to an OD_600 nm_ of 1.0. Finally, bacteria were incubated with 500 nM siderophore-^55^Fe at 30 °C. At t = 0, 15, 30, 45, 60 and 120 minutes, aliquots were removed and cells were harvested by centrifugation and the radioactivity monitored in the bacteria pellet as previously described for iron uptake by pyochelin (Hoegy and Schalk 2014). The assay was carried out as well with bacteria incubated during 15 min with 200 μM CCCP in 50 mM Tris-HCl (pH 8.0) (OD_600 nm_ of 1.0), before addition of 500 nM siderophore-^55^Fe.

### Protein expression and crystallization

Full-length *foxB* gene from *P. aeruginosa PAO1* strain was cloned into a pnEK vector with a tobacco etch virus (TEV) cleavable N-terminal His_6_-tag. Protein was expressed in *E. coli* C43 (DE3) cells grown in terrific broth medium. Cells were grown at 37 °C to an OD_600_ of 1 and induced with 0.1 mM IPTG followed by overnight growth at 20 °C.

Total membranes were prepared after cell lysis and solubilized in 1% (w/v) dodecyl maltopyranoside (DDM) in TBS buffer (20 mM Tris (pH 7.5), 350 mM NaCl, 10% glycerol) for 1 hr at 4 °C. Solubilized membranes were applied onto Ni^2+^-NTA resin and washed with TBS buffer + 0.03% DDM and 40 mM imidazole; protein was eluted with 300 mM imidazole and TEV was added overnight (1:5 w/w TEV:protein). Next day, FoxB was reverse purified to remove TEV, the cleaved His_6_-tag and any non-cleaved FoxB protein. Protein was then concentrated using a 50 kDa cut-off concentrator and injected onto a Superdex S200 10/300 increase column equilibrated with TBS buffer + 0.2 % decyl maltopyranoside (DM) without glycerol. Purified FoxB protein in TBS buffer + 0.2 % DM was concentrated to 15 mg/ml and crystallized using sitting-drop vapor diffusion.

### Ferrozine reduction assay

To check the reducing activity of FoxB, the protein was purified as described above using DDM and then incorporated into MSP1D1 nanodiscs in a 1:2:40 ratio (FoxB:MSP1D1:POPC) using previously established protocols. Briefly, proteins and lipids were mixed in the stated ratio, and biobeads (0.5 mg/ml) were added for 4 hours at 4 °C. Subsequently, assembled protein was removed from biobeads and purified further using a Superdex S200 10/300 column equilibrated in HEPES pH 7, 350 mM NaCl buffer (buffer B).

For the ferrozine assays, FoxB in nanodiscs (final concentration 3 μM) was mixed with varying concentrations of Fe^3+^-ferrioxamine B in buffer B, supplemented with 0.5 mM ferrozine and 30 mM DTT. The mixture was incubated at 37 °C for 45 min and absorbance at 562 nm was then measured. Ferrozine forms a stable magenta-colored complex with absorption peak at 562 nm with ferrous iron (Stookey 1970). As a negative control, FoxB was omitted from the reaction mixture.

### Structure determination

The structure of FoxB was determined using X-ray crystallography. Diffraction data were collected at 100 K at the PETRA III/DESY P11, P13 and P14 beamlines. Datasets were processed with XDS (Kabsch 2010) and integrated using AIMLESS (Evans 2011). Due to diffraction anisotropy StarANISO webserver (Tickle, Flensburg et al. 2018) was used to perform anisotropy correction. Heavy atom sites were calculated with SHELX C/D (Sheldrick 2008). Further density modification was carried out using RESOLVE as part of the PHENIX package (Adams, Afonine et al. 2010). A model generated by the AlphaFold2 group during the CASP14 competition (target T1058) (Callaway 2020) was used for molecular replacement and subsequent MR-SAD. Anomalous difference maps were also used to validate the model. The final model was refined using COOT (Emsley, Lohkamp et al. 2010) and REFMAC (Murshudov, Skubak et al. 2011) using thermal libation and screw-rotation (TLS) and jelly body parameters. All data collection and refinement statistics are summarized in Table S1.

## Acknowledgements

We are grateful to the staff at beamlines P11 (DESY), P13 and P14 (EMBL, Hamburg) and thank members of the Tidow lab for helpful discussions. We acknowledge access to the Sample Preparation and Characterization (SPC) Facility of EMBL. We thank the AlphaFold2 team (DeepMind/Google) for their excellent model (CASP14 target ID: T1058) and Andriy Kryshtafovych (UC Davis) for making CASP14 results available. This research was funded by the excellence cluster ‘The Hamburg Centre for Ultrafast Imaging - Structure, Dynamics and Control of Matter at the Atomic Scale’ of the Deutsche Forschungsgemeinschaft (DFG EXC 1074).

## Author Contributions

Investigation, I.J., K.V., and V.N.; Writing, I.J., I.J.S. and H.T.; Funding Acquisition & Supervision, I.J.S and H.T.

## Notes

The authors declare no competing financial interest.

## Footnotes

The abbreviations used are:

ASU: asymmetric unit
CAA: Casamino Acid
CCCP: carbonyl cyanide *m*-chlorophenylhydrazone
DDM: dodecyl maltopyranoside
DFO: Ferrioxamine B
DM: decyl maltopyranoside
CASP14: Critical Assessment of Structure Prediction 14
FoaB: ferrioxamine B
IM: inner membrane
IPTG: isopropyl β-D-1-thiogalactopyranoside
MR: molecular replacement
Ni-NTA: Ni-nitrilotriacetic acid
NOCA: nocardamine
OM: outer membrane
PCH: pyochelin
PVD: pyoverdine
SAD: single-wavelength anomalous dispersion
SEC: size-exclusion chromatography
TBDT: TonB-dependent transporter
TEV: tobacco etch virus
TM: trans-membrane

## Data Availability

Structural coordinates and structural factors have been deposited in the RCSB Protein Data Bank under accession number 7ABW (see Table S1, Suppl. Info.). All other relevant data generated during and/or analyzed during the current study are available from the corresponding author on reasonable request.

## Significance Statement

Secretion of siderophores allows most microbes to assimilate ferric ions into their biological processes. Siderophores must be taken up into the cells and chelated iron must be released. Here, we present the structure of an inner membrane siderophore reductase, FoxB, which is involved in the uptake of iron from ferrioxamine siderophores in *P.* aeruginosa. Our structure reveals FoxB to be a di-heme membrane protein, which is able to reduce the iron in chelated ferric siderophore complexes. These results offer insights into the function of this poorly characterized membrane protein family and its role in iron release from bacterial siderophores.

