## Supplementary information for "Structural insights into a novel family of integral membrane siderophore reductases"

<u>**Supplementary figures, legends and tables**</u>

**Supplementary Figure S1**

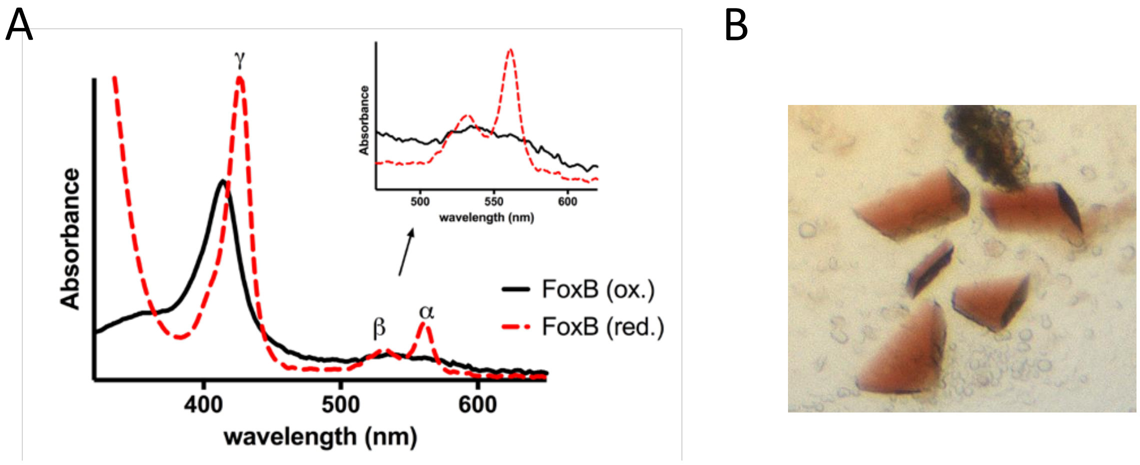

**Spectroscopic properties of heme-bound FoxB.** A) UV-Vis spectrum of FoxB showing a Soret band and a broad Q-band characteristic for heme-bound proteins. Reducing treatment of FoxB with sodium dithionite resulted in the shift of the Soret peak from 414 nm to 429 nm, along with a sharp appearance of distinct peaks in the Q-band region. Oxidation of FoxB with potassium ferricyanide reverted the spectrum to its original state, indicating that FoxB purifies in an oxidized state. B) Typical FoxB crystals showing red colour.

**Supplementary Figure S2**

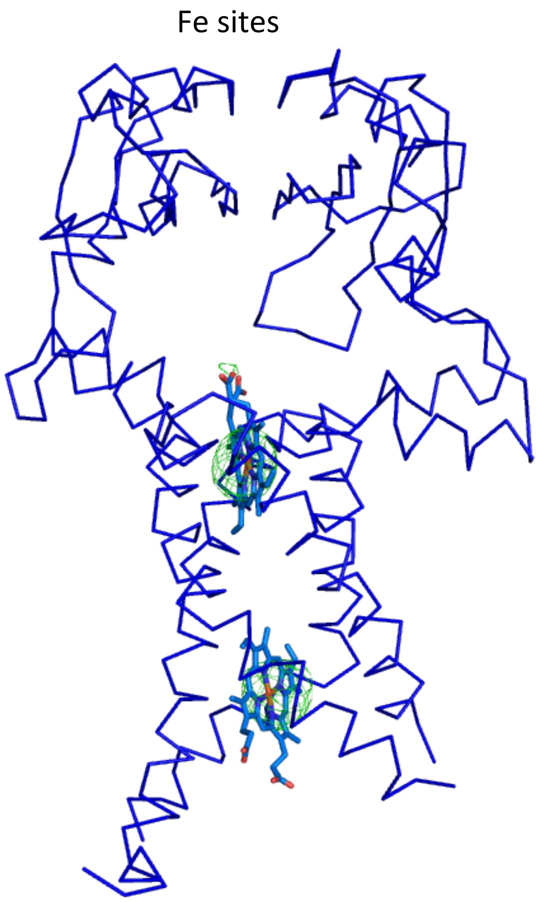

**Anomalous difference maps showing iron and selenium sites in FoxB**. Fe and Se-Met sites were determined using Phaser EP. Three out of five (including the N-terminal) methionine sites could be detected.

**Supplementary Figure S3**

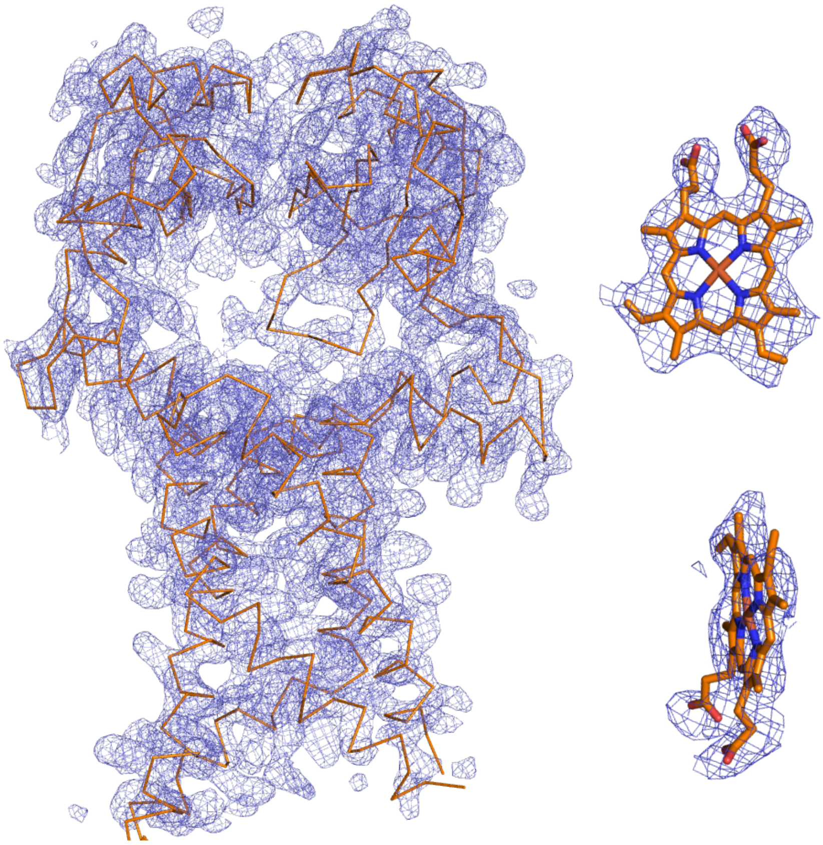

**Overall electron density for FoxB after MR-SAD phasing.** 2Fo-Fc electron density map is shown at 1.5 sigma after MR-SAD using AlphaFold2 CASP14 model and Fe/Se-Met phases.

**Supplementary Figure S4**

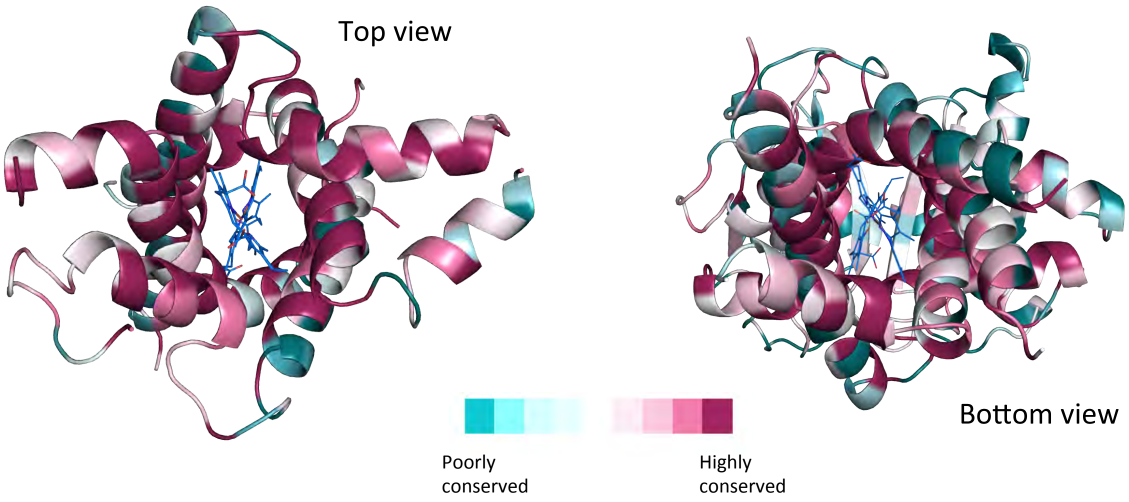

**Consurf analysis of FoxB.** The structure of FoxB was submitted to the Consurf web server to analyse the conserved regions of the model and sequence. 150 homologues with sequence conservation between 45 and 95% were automatically selected. Colour code: dark purple for highly conserved and cyan for highly variable amino acid positions.

**Supplementary Figure S5**

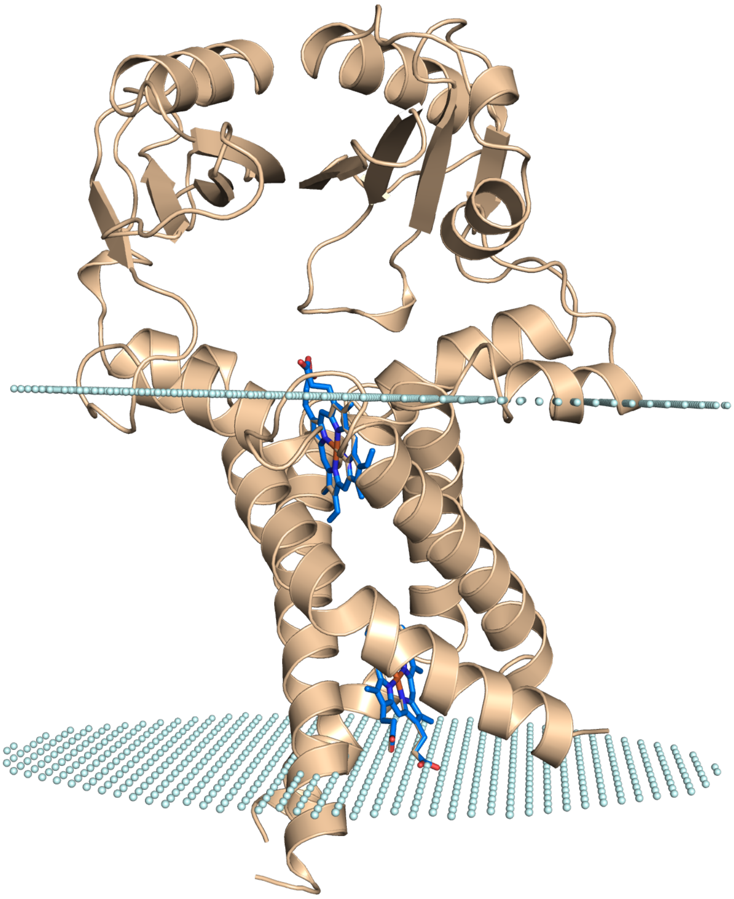

**Analysis of FoxB membrane integration.** The structure of FoxB was submitted to the OPM web server for analysis of its possible orientation within the membrane bilayer. The results show that the bilayer thickness of FoxB TM domains is 30.4 Å with a 20° tilt with respect to the bilayer.

**Supplementary Figure S6**

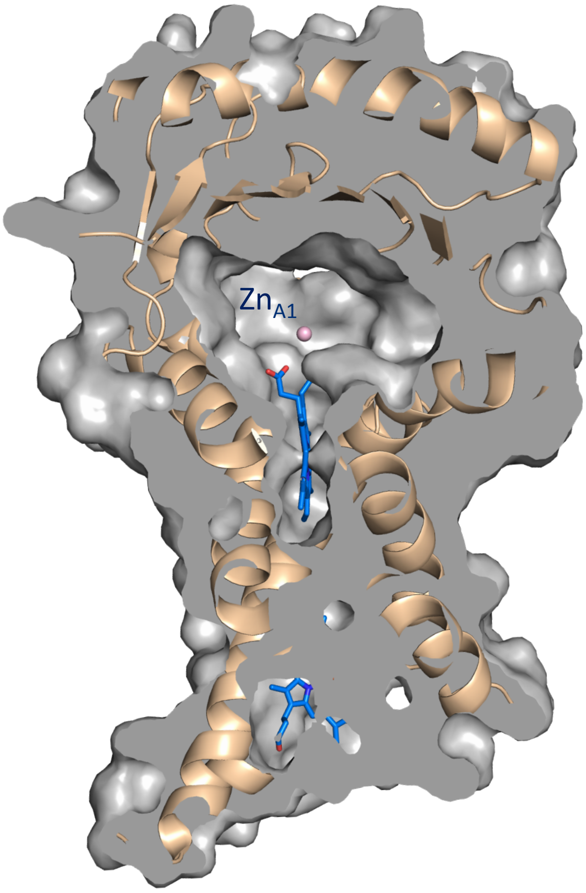

**Cut-away surface representation of FoxB showing large periplasmic cavity adjacent heme and Zn molecules.**

**Supplementary Figure S7**

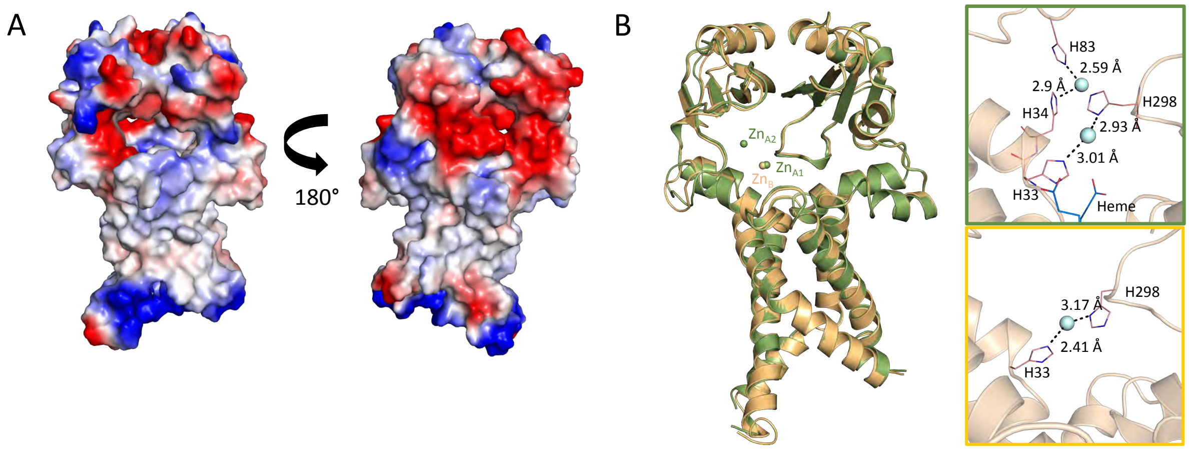

**A) Electrostatic surface potential of FoxB.** Inspection of the vacuum electrostatic charge distribution using Pymol indicates a patch of negatively charged amino acids on the surface of the periplasmic PepSY domains, which create small tunnel into the periplasmic cavity of FoxB. **B) Details of Zn coordination in periplasmic cavity.** Superposition of the two FoxB models from the crystal structure indicates that the two molecules in the asymmetric unit are almost identical. We observe Zn atoms coordinated by the FoxB molecules, which were included the crystallisation cocktail. One of the FoxB molecules has two Zn atoms bound ($Zn_{B1}$ and$Zn_{B2}$), whilst the second FoxB molecule only has one Zn atom ($Zn_{A}$). The periplasmic PepSY domains superimpose with rmsd values of > 3 Å with other bacterial proteins containing these domains. The function of PepSY domains in these proteins remains to be determined.

**Supplementary Figure S8**

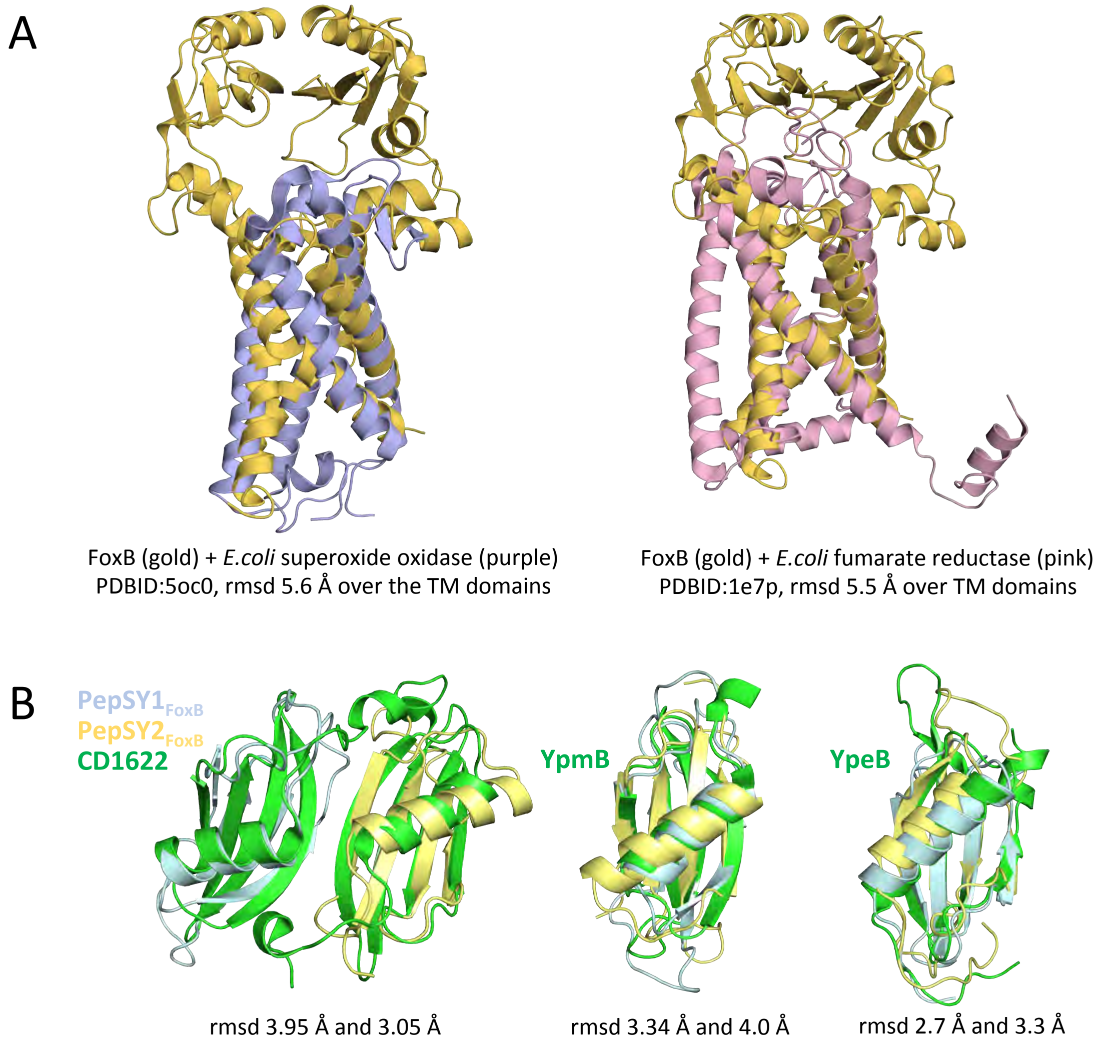

**Structural homologs of FoxB.** A) Structural homologs of FoxB transmembrane region as identified by DALI server. DALI server identified two proteins *E. coli* superoxide oxidase and fumarate reductase, which exhibit a distant structural resemblance to the TM domains for FoxB (rmsd > 5.5 Å). B) Superposition of FoxB PepSY domains with homologous domains. While in general all PepSY domains superimpose well, none of them could be used successfully for molecular replacement.

**Supplementary Figure S9**

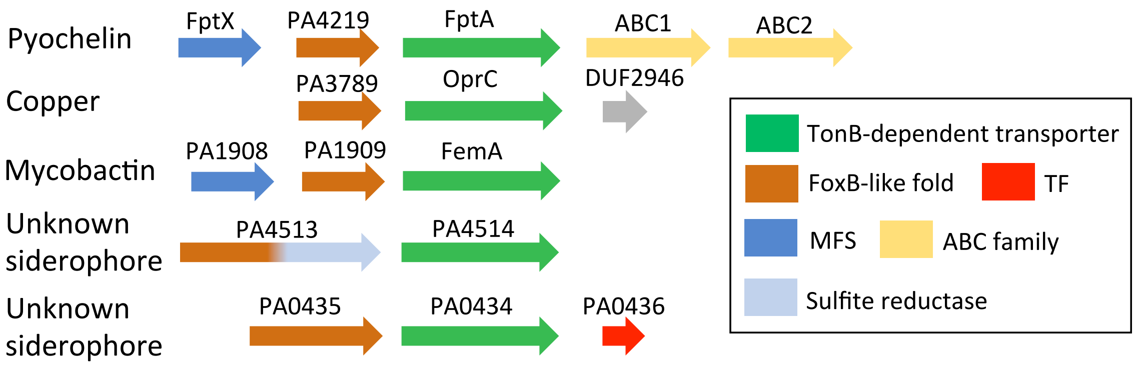

**Additional siderophore/metal uptake operons in *P. aeruginosa* containing members of the FoxB (COG3182) family.** Genes containing a similar topology with 4 TM helices and PepSY domains, belonging to the COG3182 family of membrane proteins were further identified in *P. aeruginosa* using BLAST with the sequence of FoxB. The ferrioxamine and pyoverdine operons are omitted from the representation. PA0453 is predicted to contain an extra TM helix.

**Table S1: Data collection and refinement statistics**

| | FoxB dataset* (PDBID: 7ABW) |
|---|---|
| **Data collection** | |
| **Beamline** | PETRA III, P11 |
| Space group | $P22_12_1$ |
| Cell dimensions | |
| *a*, *b*, *c* (Å) | 64.4, 114.3, 237.1 |
| α, β, γ (°) | 90, 90, 90 |
| Resolution (Å) | 46.3-3.07 (3.4-3.07) |
| $R_{\mathrm{pim}}$ | 0.025 (0.99) |
| $R_{meas}$ | 0.079 (2.77) |
| $I / \sigma I$ | 18.9 (1.1) |
| $CC_{1/2}$ | 0.99 (0.39) |
| Completeness (%)<br>spherical<br>ellipsoidal | <br>60.5 (11.4)<br>92.5 (73.3) |
| Redundancy | 19.1 (13.3) |
| **Refinement** | |
| Resolution (Å) | 3.074 |
| No. reflections | 20228 (1011) |
| $R_{\mathrm{work}}$ / $R_{\mathrm{free}}$ | 0.275/0.299 |
| No. atoms | 5949 |
| Protein | 5860 |
| Ligand/ion | 89 |
| *B*-factors | |
| Protein | 49.8 |
| Ligand/ion | 42.66 |
| R.m.s. deviations | |
| Bond lengths (Å) | 0.0145 |
| Bond angles (°) | 2.4 |
| Ramachandran (%)<br>Favored regions<br>Allowed regions<br>Outliers | <br>91.33<br>7.83<br>0.84 |

*Values in parentheses are for highest-resolution shell.

### Table S2. Strains used in this study

| Strain | Collection ID | Relevant characteristics | Source or references |
|---|---|---|---|
| ***Escherichia coli*** | | | |
| TOP10 | | F-, *mcrA* Δ(*mrr-hsdRMS-mcrBC*), ϕ80*lacZ*, Δ*M15*, Δ*lacX74*, *recA1*, *araD139*, Δ(*ara-leu*)7697, *galU*, *galK*, *rpsL*, *endA1*, *mup* | Invitrogen |
| ***Pseudomonas aeruginosa*** | | | |
| PAO1 | PAO1 | Wild-type strain | (Stover, Pham et al. 2000) |
| *ΔpvdFΔpchA* | PAS283 | PAO1; *pvdF* and *pchA* chromosomally deleted | (Gasser, Baco et al. 2016) |
| *ΔpvdFΔpchAΔfoxA* | PAS535 | PAO1; *pvdF*, *pchA* and *foxA* chromosomally deleted | (Normant, Josts et al. 2020) |
| *ΔpvdFΔpchAΔfoxB* | PAS600 | PAO1; *pvdF*, *pchA* and *foxB* chromosomally deleted | This study |
| ***Plasmids*** | | | |
| pEXG2 | pEXG2 | allelic exchange vector with pBR origin, gentamicin resistance, sacB | (Rietsch, Vallet-Gely et al. 2005) |
| pEXG2Δ*foxB* | pVN9 | pEXG2 carrying the sequence to delete *foxB* | This study |

### Table S3. Primers used for RT-qPCR analysis

| Name | Gene | Sequence |
|---|---|---|
| uvrD F | *uvrD* | CTACGGTAGCGAGACCTACAACAA |
| *uvrD* R | *uvrD* | GCGGCTGACGGTATTGGA |
| *foxA* F | *foxA* | AAGGGCTCGGATACCCAGTT |
| *foxA* R | *foxA* | CGTTGGGATCGTGTTGCA |
| *foxB* F | *foxB* | GATGTTCTGGTTCCTCGACTG |
| *foxB* R | *foxB* | TTGCCGCGCTTGATCTT |

### Table S4. Primers used for the construction of the *ΔpvdF ΔpchA ΔfoxB* strain

| Name | Gene | Sequence |
|---|---|---|
| FoxB XhoI F | *foxB* | AAAATCTCGAGCCGGCACGCCCCTGGCGCCCACCGAGGGCAAGCAG TGG |
| FoxB overlap F | *foxB* | GGTTACCTGGAGCCCCCGGTGGCGCAAGCGCCGGGCGCGCCACTGG |
| FoxB overlap R 2 | *foxB* | GGGAGGCAGCCAGACTAACAGCCGGGGGGCTCCAGGTAACCAGGGT ACGGCCGGCGACG |
| FoxB HindIII R 2 | *foxB* | AAAACAAGCTTCGCTCGTGGCCCGGCAGAAAGAAGCCCCGCCAAGG |
